# Reduction in myocardial fibrosis in MDX mice on oral consumption of *Aureobasidium Pullulans* produced Neu REFIX Beta-glucans; holds potential as an adjuvant in managing post-transplantation organ fibrosis

**DOI:** 10.1101/2023.09.25.559276

**Authors:** Senthilkumar Preethy, Naoki Yamamoto, Kottoorathu Mammen Cherian, Rajasekaran Premsekar, Gary A Levy, Rajappa Senthilkumar, Samuel JK Abraham

**Affiliations:** Fujio-Eiji Academic Terrain (FEAT), Nichi-In Centre for Regenerative Medicine (NCRM), Chennai, India; National Centre for Global health and Medicine (NCGM), Chiba, Japan; Frontier Lifeline Hospitals, R-30-C Ambattur Industrial Estate Road, Mogappair, Chennai - 600 101, Tamilnadu, India; Dr. Kamakshi Memorial Hospital, Chennai, India; Emeritus professor, Medicine and Immunology, University of Toronto, Ontario, Canada; Antony- Xavier Interdisciplinary Scholastics (AXIS), GN Corporation Co. Ltd., Kofu, Japan; Centre for Advancing Clinical Research (CACR), University of Yamanashi - School of Medicine, Chuo, Japan; Mary-Yoshio Translational Hexagon (MYTH), Nichi-In Centre for Regenerative Medicine (NCRM), Chennai, India; R & D, Sophy Inc., Japan; Levy-Jurgen Transdisciplinary Exploratory (LJTE), Global Niche Corp, Wilmington, DE, USA

**Keywords:** Beta-glucan, Duchenne Muscular Dystrophy (DMD), Organ fibrosis, Transplantation, Collage Type I

## Abstract

Organ fibrosis is one of the major causes of morbidity and mortality globally. Though fibrosis in genetic diseases such as Duchenne muscular dystrophy (DMD) may be attributed to the genetic defect, chronic microinflammation remains a key mechanism underlying such fibrosis, which also precedes both other organ fibrosis and post-organ transplant fibrosis. Having proven the anti-inflammatory, anti-fibrotic effects of Beta-1,3-1,6-glucan (Neu-REFIX) produced by N-163 strain of *Aureobasidium Pullulans* in earlier clinical and pre-clinical studies, we performed the current study to evaluate its effects on myocardial fibrosis. N-163 beta-glucan was administered to 45 mice in three groups, each fifteen animals, Gr. 1, normal mice, Gr.2, mdx mice as vehicle, Gr.3, mdx mice which were administered Neu REFIX beta-glucan orally. Evaluation of Collagen Type I (Col-I) in myocardium was performed by immunohistochemistry. Percentage of myocardium Col-I positive area of 6.42 ± 2.67 significantly decreased in the Neu-REFIX group (4.32 ± 1.78) (p-value < 0.01). As myocardial fibrosis has been shown to be reduced following treatment with N-163 beta glucan in a genetic, muscle structure anomaly disease such as DMD, in addition to adding value to DMD patients, in whom myocardial failure occurs in the advanced stages leading to pre-mature death, Neu-REFIX beta-glucan adjuvant treatment in the setting of solid organ transplantation may be of value to reduce the incidence of fibrosis which is a known feature of chronic allograft rejection leading to graft loss.

## Introduction

Duchenne muscular dystrophy (DMD) is a hereditary muscle condition caused by mutations in the DMD gene on the X chromosome, which codes for the dystrophin protein. This dystrophin deficiency in myofibers causes contraction-induced membrane damage with release of cytoplasmic contents and stimulation of innate immunity, cycles of myofiber degeneration/regeneration, age-related muscle replacement by fibrofatty connective tissue, muscle weakness, and, eventually, death. Dystrophin deficiency causes chronic activation of the immune system and associated chronic inflammatory response [1]. In contrast to a single bout of inflammation and repair of normal muscle, the dystrophin-deficient muscle loses this bout effect owing to the chronic inflammatory state. Asynchronous regeneration results in skeletal fibrosis which is followed by fibrosis of other muscles including the cardiac muscle. Myocardial fibrosis progresses to dilated cardiomyopathy, which is worsened by heart failure and arrhythmia [2]. Even while recent advances in the therapy of respiratory insufficiency have increased the longevity and overall prognosis of DMD patients, unexpected fatalities due to heart failure have a severe impact on their quality of life [3]. Prompt treatment and early identification of cardiomyopathy are prerequisites for effective cardioprotective treatments that prevent or reduce the processes of ventricular remodelling and heart failure. While there is no definitive cure for the disease, approaches to control inflammation by use of corticosteroids has been a standard of care but they have their own associated side effects [4]. Medications that target fibrosis including cardiac fibrosis are in the early stages of development. We have previously reported the anti-inflammatory, anti-fibrotic and immune-modulating effects of a biological response modifier beta-glucan (BRMG) produced by the N-163 strain of *Aureobasidium pullulans* (*A*.*pullulans*) (Trade name: Neu-REFIX) in pre-clinical mdx mice model [5,6] and human clinical studies of DMD [7,8]. In the current study we wanted to evaluate the effects of this Neu-REFIX beta-glucan food supplement on myocardial fibrosis in mdx mice model of DMD.

## Methods

The study comprised 45 mice [3], who were randomised into three groups of fifteen mice each the day before the commencement of therapy based on their body weight. Using Excel software, body weight-stratified random sampling was used for randomization.

Group 1: Normal-Fifteen C57BL/10SnSlc mice were without any treatment until sacrifice.

Group 2: Vehicle-Fifteen mdx mice were orally administered vehicle [pure water] at a volume of 10 mL/kg once daily from days 0 to 45.

Group 3: N-163 beta-glucan -Fifteen mdx mice were orally administered with N-163 strain produced B-glucan (Neu-REFIX™), was provided by GN Corporation Co. Ltd at a dose of 3 mg/kg as API in a volume of 10 mL/kg once daily for 45 days.

On day 45, the animals were sacrificed through abdominal vena cava exsanguination under isoflurane anaesthesia (Pfizer Inc.). The myocardium muscles were collected and measured. For immunohistochemistry, the frozen muscle sections (myocardium, 6 mice/group) were cut from frozen blocks and fixed in acetone. Endogenous peroxidase activity was blocked using 0.03% H2O2 for 5 minutes, followed by incubation with Block Ace (Dainippon Sumitomo Pharma Co. Ltd., Japan) for 10 minutes. The sections were incubated with primary antibody at 4°C for overnight. After incubation with secondary antibody, enzyme substrate reactions were performed using 3, 3’-diaminobenzidine/H2O2 solution (Nichirei Bioscience Inc., Japan). Profiles of primary and secondary antibodies are shown in Table 1.

**Table 1.**
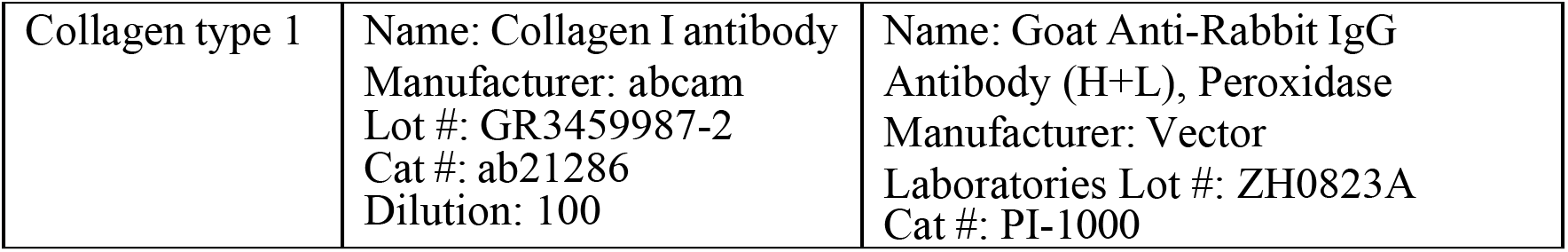

For quantitative analysis of collagen type 1-positive area, bright field images of collagen type 1-immunostained sections were captured using a digital camera (DFC295; Leica) at 200-fold magnification, and the positive areas in 5 fields/section were measured using ImageJ software (National Institute of Health).

Statistical analyses were performed using Prism Software 6 (GraphPad Software, USA). Statistical analyses were performed using Bonferroni Multiple Comparison Test. Comparisons were made between the following groups;

1. Group 2 (Vehicle) vs. Group 1 (Normal), Group 3 (Neu-REFIX beta-glucan). P values <0.05 was considered statistically significant. Results were expressed as mean ± SD.

A trend or tendency was assumed when a one-sided t-test returned P values <0.1. Comparisons were made between the following groups;

1. Group 2 (Vehicle) vs. Group 1 (Normal)
2. Group 2 (Vehicle) vs. Group 3 (Neu-REFIX beta-glucan)

## Results

The mean body weight of mice in the vehicle group was significantly lower than that of mice in the normal group but there was no significant difference in the mean body weight between the vehicle group and Neu-REFIX beta-glucan treated mice. Representative photomicrographs of the Col-I-immunostained sections are shown in Figure 1. There was an increase in the percentage of Col-I positive area in the vehicle group (6.42± 2.67) compared to the normal mice (1.72 ± 0.78) while it significantly decreased in the Neu-REFIX group (4.32 ± 1.78) (p-value < 0.01). (Figure 2)

**Figure 1.**
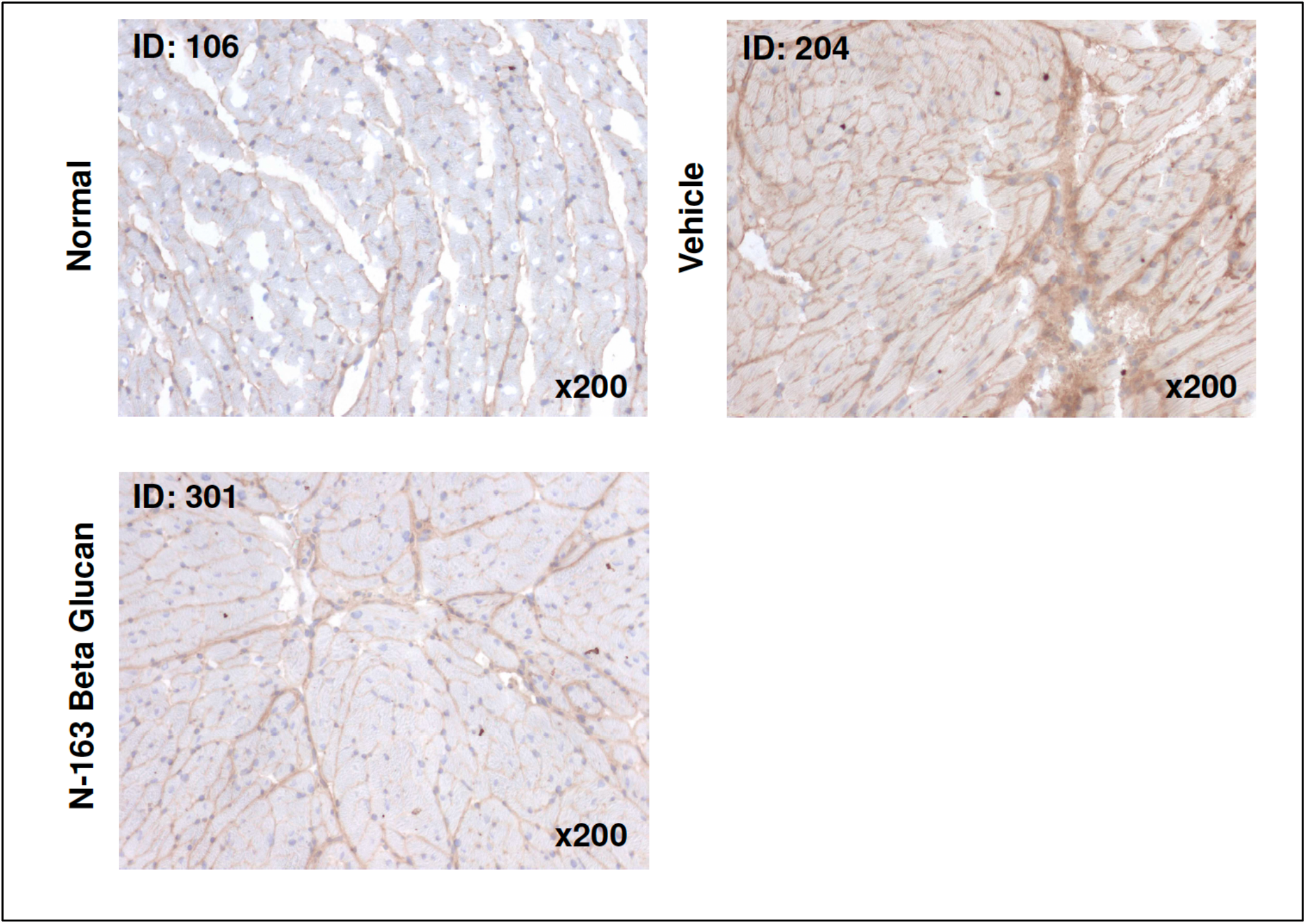
Representative photomicrographs of collagen type 1-immunostained muscle sections (myocardium) showing a marked reduction in collagen deposition.

**Figure 2.**
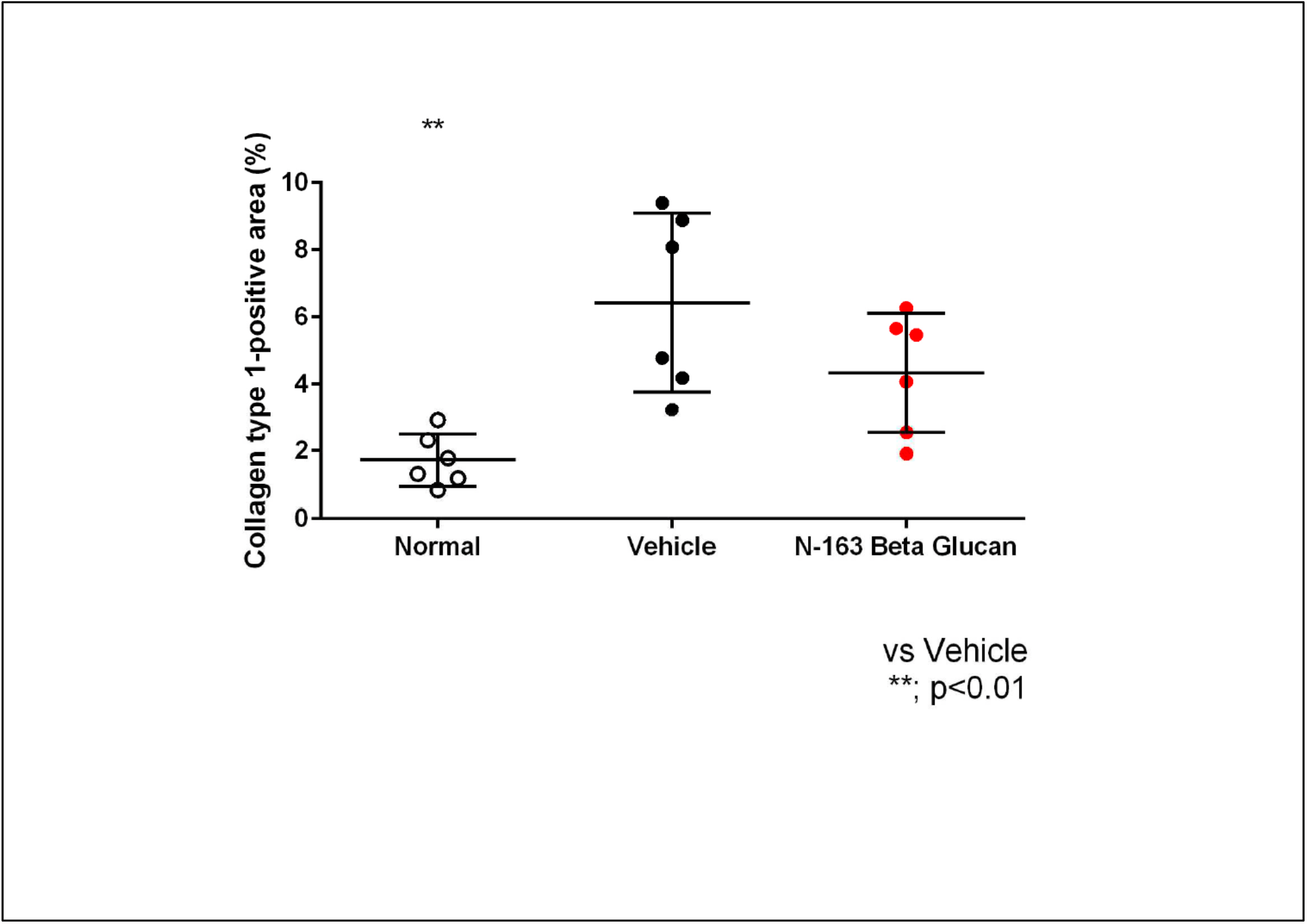
Collagen type 1-positive area at 200-fold magnification, and the positive areas in 5 fields/section. Data is the mean and SD from six mice; p-value significance between Group 2 (Vehicle) vs. Group 3 (N-163 beta-glucan)

## Discussion

Fibrosis is one of the early clinical markers of dystrophic cardiac disease, appearing in DMD patient hearts before the age of 10 years. Cardiomyocytes, fibroblasts, endothelial cells, and immune cells are all involved in the formation of fibrosis in the heart. In response to stimuli such as inflammation, mechanical signals, and signals from neighbouring cells, all these cells can generate profibrotic cytokines and chemokines [9]. Cellular stress and necrotic death in cardiomyocytes most likely initiate fibrosis in the dystrophic heart by infiltrating the surrounding tissue with cytokines, chemokines, and debris that attract neutrophils and macrophages. Fibrosis likely causes additional functional myocardial loss by increasing the workload requirement on neighbouring cardiomyocytes, which will now be required to do excessive contractile effort in a stiff, mechanically stressful environment. Second, the fibrotic scars may disrupt electrical conduction channels across the heart, potentially leading to a variety of arrhythmias [10]. Because the fibrotic process begins so early in the disease’s progression and the effects of untreated fibrosis can be so severe, decreasing fibrosis has been a significant focus of cardiac-directed DMD therapy. Angiotensin-converting enzyme (ACE) inhibitors, angiotensin II receptor blockers (ARBs), mineralocorticoid receptor antagonists, statins such as Simvastatin and anti-diabetic drugs such as Metformin have been employed in studies to prevent the build-up of fibrosis in dystrophic animal and human hearts but not with significant clinical outcome [2,10, 11].

Although solid organ transplantation is now recognized as the best treatment for patients with end stage organ failure, long term survival rates of solid organ grafts have not improved significantly and within 10 years post-transplant 40-50% are lost due to both immunologic and non-immunologic factors. In the setting of heart transplantation, both ischemia-reperfusion injury (IRI) and chronic allograft rejection, are major factors contributing to chronic rejection and fibrosis is hallmark of chronic rejection. When cardiomyocytes are damaged, the process results in production of collagen leading to fibrosis. Damage to endothelial cells (EC) leads to reduced myocardial perfusion resulting in further hypoxia in affected areas. Hypoxia, in turn, has been shown to induce phenotypic changes consistent with endothelial-to-mesenchymal transition (EndMT), in which ECs undergo phenotypic changes and transdifferentiate into myofibroblast-like cells with increased extracellular matrix protein production, contributing to fibrosis [12]. Also, by secreting cytokines, adaptive immune cells (such as B and CD4+ T cells) and innate immunity cells (such as neutrophils and innate lymphoid cells) contribute to the trans differentiation of recipient-derived macrophages to myofibroblasts, which leads to transplanted organ fibrosis [13]. Although the mechanisms leading to chronic allograft rejection including the role of innate immune mechanisms including macrophage and NK cell activation and antibody mediated rejection are better understood, there has been little progress in the treatment of chronic allograft rejection.

Corticosteroids, remain the only effective pharmacotherapy for DMD, extend independent ambulation by 2 to 4 years but have problematic side effects. Although prednisone was first studied to reduce muscular inflammation, its therapeutic mechanisms in DMD remain unknown. There is currently no effective medicine to reduce muscular fibrosis in DMD patients. Targeting fibrogenic cytokines such as TGF-Beta, suppressing muscle inflammation, and enhancing muscle regeneration are proposed as potential therapies for fibrosis in DMD, but a combination of antifibrotic therapies targeting different aspects of muscle fibrogenesis is likely to be required [14]. In the case of post-transplant fibrosis too, anti-fibrotic medicines that target the fibrosis at several levels, including epigenetic enzymes, genes, translation machinery, and signalling molecules are being experimented. Despite encouraging pre-clinical outcomes, there are several obstacles that might jeopardise the clinical translation of such anti-fibrotic therapies [15,16]

In both DMD and post-transplant fibrosis, targeting the immune-inflammatory response has been advocated as a potential therapeutic strategy [1, 15]. Additionally, if the therapeutic strategy can also address inflammation and enhance muscle regeneration, it will be ideal. The N-163 beta-glucan (Neu-REFIX) strategy has demonstrated its potential in reducing DMD fibrosis and inflammation in earlier clinical and pre-clinical studies by decreasing the inflammation score, plasma TGF-β levels and plasma IL-13 apart from enhancing muscle regeneration and clinically relevant improvement in ambulation [5-8]. In the current study, its effects on significant decrease in collagen Type-I in the myocardium of mdx mice proves its potential as a anti-fibrotic agent in both DMD as well as myocardial fibrosis at a tissue level.

The limitations of the study include that it is only a pre-clinical study and the molecular mechanisms of pathogenesis and contributing factors being more complex and diverse in clinical settings of DMD and post-transplant fibrosis, long term clinical studies are mandatory to validate the potential of Neu-REFIX beta-glucan as a potential pharmacological adjuvant in DMD and organ-transplantation associated fibrosis

## Conclusion

This study has confirmed the safety of oral consumption of Neu-REFIX beta 1.3-1,6 glucans in reducing the myocardial fibrosis, in this mdx model of DMD which is a genetic disease animal model. These encouraging results unravel the potentials of this beta glucans, worth trying in DMD patients as an orally administrable drug adjuvant in the prevention and management of myocardial fibrosis and also, other organ fibrosis, post-organ transplant fibrosis in larger clinical studies.

## Acknowledgments

The authors thank,

- Dr. Yoshitsugu Aoki and his colleagues of National Center of Neurology and Psychiatry (NCNP), Japan, for their technical guidance with the handling of tissue specimens and their preservation, that were used in this study.
- Mr. Yasushi Onaka and Mr. Masato Onaka, Ms Misa Takamoto of Sophy Inc. for technical clarifications.
- Ms. Eiko Amemiya of II Dept. of Surgery, University of Yamanashi for secretarial assistance.
- Ms. Yoshiko Amikura of GN Corporation, Japan for liaising between the institutes.

## References

1. Farini A, Gowran A, Bella P, Sitzia C, Scopece A, Castiglioni E, Rovina D, Nigro P, Villa C, Fortunato F, Comi GP, Milano G, Pompilio G, Torrente Y. Fibrosis Rescue Improves Cardiac Function in Dystrophin-Deficient Mice and Duchenne Patient-Specific Cardiomyocytes by Immunoproteasome Modulation. Am J Pathol. 2019 Feb;189(2):339–353.

2. Meyers TA, Townsend D. Cardiac Pathophysiology and the Future of Cardiac Therapies in Duchenne Muscular Dystrophy. Int J Mol Sci. 2019 Aug 22;20(17):4098.

3. Schultz TI, Raucci FJ Jr, Salloum FN. Cardiovascular Disease in Duchenne Muscular Dystrophy: Overview and Insight Into Novel Therapeutic Targets. JACC Basic Transl Sci. 2022 Mar 9;7(6):608–625.

4. Gloss D, Moxley RT 3rd, Ashwal S, Oskoui M. Practice guideline update summary: Corticosteroid treatment of Duchenne muscular dystrophy: Report of the Guideline Development Subcommittee of the American Academy of Neurology. Neurology. 2016 Feb 2;86(5):465–72.

5. Preethy S, Sakamoto S, Higuchi T, Ichiyama K, Yamamoto N, Ikewaki N, Iwasaki M, Dedeepiya VD, Srinivasan S, Rajmohan M, Senthilkumar R, Abraham S. Enhanced muscle regeneration in mdx mice, Duchenne muscular dystrophy animal model, proven by CD44 & MYH3 expression, on oral feeding of N-163 strain of Aureobasidium Pullulans produced B-Glucan. biorxiv 2023.06.06.543858v1 doi: 10.1101/2023.06.06.543858

6. Preethy S, Aoki Y, Minegishi K, Senthilkumar R, Abraham S. Resolution of fibrosis in mdx dystrophic mouse after oral consumption of N-163 strain of Aureobasidium pullulans produced biological response modifier β-glucan (BRMG). bioRxiv 2022.11.17.516628; doi: 10.1101/2022.11.17.516628

7. Raghavan K, Dedeepiya VD, Srinivasan S, Pushkala S, Subramanian S, Ikewaki N, Iwasaki M, Senthilkumar R, Preethy S, Abraham S. Beneficial immune-modulatory effects of the N-163 strain of Aureobasidium pullulans-produced 1,3-1,6 Beta glucans in Duchenne muscular dystrophy: Results of an open-label, prospective, exploratory case-control clinical study. IBRO Neuroscience reports 2023; 15: 90–99. doi: 10.1016/j.ibneur.2023.06.007.

8. Raghavan K, Sivakumar T, Bharatidasan SS, Srinivasan S, Dedeepiya VD, Ikewaki N, Senthilkumar R, Preethy S, Abraham S. Efficacy of N-163 strain of Aureobasidium pullulans-produced beta-glucan in improving muscle strength and function in patients with Duchenne muscular dystrophy; Results of a 6-month non-randomised open-label linear clinical trial. medRxiv 2023.04.29.23289260v1; doi: 10.1101/2023.04.29.23289260

9. Tandon A, Villa CR, Hor KN, Jefferies JL, Gao Z, Towbin JA, Wong BL, Mazur W, Fleck RJ, Sticka JJ, Benson DW, Taylor MD. Myocardial fibrosis burden predicts left ventricular ejection fraction and is associated with age and steroid treatment duration in duchenne muscular dystrophy. J Am Heart Assoc. 2015 Mar 26;4(4):e001338.

10. Ma Y, Mouton AJ, Lindsey ML. Cardiac macrophage biology in the steady-state heart, the aging heart, and following myocardial infarction. Transl Res. 2018 Jan;191:15–28. doi: 10.1016/j.trsl.2017.10.001

11. Angebault C, Panel M, Lacôte M, Rieusset J, Lacampagne A, Fauconnier J. Metformin Reverses the Enhanced Myocardial SR/ER-Mitochondria Interaction and Impaired Complex I-Driven Respiration in Dystrophin-Deficient Mice. Front Cell Dev Biol. 2021 Jan 25;8:609493.

12. Hurskainen M, Ainasoja O, Lemström KB. Failing Heart Transplants and Rejection-A Cellular Perspective. J Cardiovasc Dev Dis. 2021 Dec 12;8(12):180. doi: 10.3390/jcdd8120180.

13. Li X, Wu J, Zhu S, Wei Q, Wang L, Chen J. Intragraft immune cells: accomplices or antagonists of recipient-derived macrophages in allograft fibrosis? Cell Mol Life Sci. 2023 Jul 3;80(7):195. doi: 10.1007/s00018-023-04846-0.

14. Zhou L, Lu H. Targeting fibrosis in Duchenne muscular dystrophy. J Neuropathol Exp Neurol. 2010 Aug;69(8):771–6.

15. Toldo S, Quader M, Salloum FN, Mezzaroma E, Abbate A. Targeting the Innate Immune Response to Improve Cardiac Graft Recovery after Heart Transplantation: Implications for the Donation after Cardiac Death. Int J Mol Sci. 2016 Jun 17;17(6):)

16. Raziyeva K, Kim Y, Zharkinbekov Z, Temirkhanova K, Saparov A. Novel Therapies for the Treatment of Cardiac Fibrosis Following Myocardial Infarction. Biomedicines. 2022 Sep 2;10(9):2178.

